# Ablation of NMDA receptors in dopamine neurons disrupts attribution of incentive salience to reward-paired stimuli

**DOI:** 10.1101/439745

**Authors:** Przemysław Eligiusz Cieślak, Jan Rodriguez Parkitna

## Abstract

Midbrain dopamine (DA) neurons play a crucial role in the formation of conditioned associations between environmental cues and appetitive events. Activation of N-methyl-D-aspartate (NMDA) receptors is a key mechanism responsible for the generation of conditioned responses of DA neurons to reward cues. Here, we tested the effects of the cell type-specific inactivation of NMDA receptors in DA neurons in adult mice on stimulus-reward learning. Animals were trained in a Pavlovian learning paradigm in which they had to learn the predictive value of two conditioned stimuli, one of which (CS+) was paired with the delivery of a water reward. Over the course of conditioning, mutant mice learned that the CS+ predicted reward availability, and they approached the reward receptacle more frequently during CS+ trials than CS− trials. However, conditioned responses to the CS+ were weaker in the mutant mice, possibly indicating that they did not attribute incentive salience to the CS+. To further assess whether the attribution of incentive salience was impaired by the mutation, animals were tested in a conditioned reinforcement test. The test revealed that mutant mice made fewer instrumental responses paired with CS+ presentation, confirming that the CS+ had a weaker incentive value. Taken together, these results indicate that reward prediction learning does occur in the absence of NMDA receptors in DA neurons, but the ability of reward-paired cues to invigorate and reinforce behavior is lost.

---

The ability to use environmental stimuli to predict rewards is essential for adaptive behavior. During stimulus-reward learning, an initially neutral environmental stimulus (conditioned stimulus, CS) that is repeatedly paired with reward delivery (unconditioned stimulus, US) acquires predictive and incentive motivational properties that are used to predict reward availability and drive reward-seeking behavior. The midbrain dopamine (DA) neurons, originating in the ventral tegmental area (VTA) and substantia nigra, have been proposed to be crucial for the formation of conditioned associations between environmental cues and appetitive events. It was observed that during stimulus-reward learning, there is a transition from slow tonic to rapid phasic firing in midbrain DA neurons, causing transient phasic DA release in terminal regions, encoding reward prediction errors and attributing incentive salience to reward cues **[1–5]**. Thus, phasic DA signaling may serve as a neural substrate for both reward prediction and incentive learning. In line with this, it was shown that phasic stimulation of DA neurons was sufficient to drive behavioral conditioning and increase the willingness to engage in reward-seeking behavior **[6–9]**.

The cellular mechanism enabling reward cues to produce conditioned responses in DA neurons necessitates synaptic strengthening of excitatory inputs onto DA neurons **[10, 11]**. The key mechanism responsible for the generation of phasic activity and synaptic strengthening of excitatory synapses in DA neurons is the activation of N-methyl-D-aspartate (NMDA) receptors **[10, 11, 12–14]**. It was shown that NMDA receptor antagonism in the VTA was sufficient to block the acquisition of cue-reward associations **[10]**. However, studies performed on animals with DA neuron-specific genetic inactivation of NMDA receptors may contradict these observations. While targeted mutation in DA neurons prevented cocaine-induced strengthening of excitatory synapses and robustly decreased phasic DA release in response to food-paired cues, mutant animals were still able to develop conditioned behavioral responses to reward-paired stimuli **[15–17]**. This calls into question the role of NMDA receptor-dependent signaling in DA neurons as a key mechanism underlying stimulus-reward learning. As a way to reconcile these observations, it was proposed that the preserved learning ability could be due to compensatory mechanisms, such as increased AMPA receptor-mediated excitatory postsynaptic currents, that develop if the loss of NMDA receptors occurs during early development. These adaptations can be avoided if inactivation of NMDA receptors occurs after development is complete. It was shown that inducible inactivation of NMDA receptors in adult animals causes a complete loss of NMDA receptor-dependent phasic firing in midbrain DA neurons but does not lead to compensatory changes in synaptic transmission **[15, 18]**, suggesting that an inducible system for mutagenesis in DA neurons might be more suitable for testing a functional role of NMDA receptor-dependent signaling in reward learning.

Here, to determine the role of NMDA receptor-dependent signaling in DA neurons in stimulus-reward learning, we used NR1^DATCreERT2^ mice in which a cell type-specific inactivation of NMDA receptors can be induced in adulthood. NR1^DATCreERT2^ mice harbor a tamoxifen-inducible Cre recombinase expressed under the control of the DA transporter (DAT) promoter and a floxed variant of the *Grin1* gene encoding the NR1 subunit. As a result of the induction of mutation, no functional NMDA receptors are formed in DA neurons. The generation, breeding, genotyping and validation of these mice were described previously **[15, 18]**. A cohort of male NR1^DATCreERT2^ mice (Cre Tg/0; *Grin1* flox/flox; n = 12) and control animals (Cre 0/0; *Grin1* flox/flox or Cre 0/0; *Grin1* flox/wt; n = 12) were used in the study. Animals were housed 2-5 per cage in a room with a controlled temperature set at 22 ± 2°C under a 12 h light/dark cycle. Mice had *ad libitum* access to standard rodent laboratory chow but access to water was limited to 1-1.5 ml per day. This water restriction was maintained for the duration of the behavioral experiments. Experiments were performed in 6 standard mouse operant chambers (ENV-307-CT, Med Associates Inc.) enclosed in cubicles fitted with a fan to provide ventilation and mask extraneous noise. Each chamber was equipped with a liquid receptacle located at the center of the front wall. Saccharin-flavored water (0.01% w/v saccharin, Sigma-Aldrich), used as an US, was delivered into a receptacle by syringe pump (PHM-100, Med Associates Inc.). To avoid auditory or vibration signals associated with US delivery, syringe pumps were placed outside the sound-isolating enclosure and were connected to the liquid receptacle via a silicone tube. A house light and tone generator (65 dB, 2.9 kHz) were located on the wall opposite the liquid receptacle. All procedures were carried out in accordance with EU Directive 2010/63/EU.

First, mice were familiarized with the operant chambers and the liquid reward for two consecutive days. During adaptation, 20 μl of water was delivered into the receptacle at 60 s intervals (30 US presentations in 30 minutes). Both the control and NR1^DATCreERT2^ animals showed an increase in the number of receptacle entries over consecutive days of adaptation (**Figure 1**, main effect of session, *p* < 0.0001; complete results of statistical analyses are listed in **Table 1**). The average number of receptacle entries did not differ between genotypes, indicating that the initial willingness to engage in reward-seeking behavior was not affected by the mutation. Then, animals received 11 conditioning sessions (once daily) following the procedure described by O’Connor and colleagues **[19]**, with modifications. Each session consisted of 20 trials in which presentation of a stimulus was paired with delivery of 20 μl of water (CS+) and 20 trials in which presentation of an alternative stimulus was not paired with a reward (CS−). A 10 s constant tone and 10 s flashing (1 Hz) house light served as conditioned stimuli (counterbalanced between animals). CS+ and CS− trials were presented randomly and were separated by variable intertrial intervals (range: 30-90 s, mean: 60 s). The probability of making an entry into the water receptacle during CS+ trials significantly increased over the course of conditioning, while the probability of entering a receptacle during CS− trials remained at a constant level, suggesting that both groups of mice were sensitive to CS-US contingency (**Figure 2A**, main effect of stimulus and stimulus × session interaction, both *p* < 0.0001). The water reward was delivered 5 s after CS+ onset. To further determine if animals learned the predictive value of the reward-paired CS, we calculated the number of receptacle entries before reward delivery (discriminated approaches). The analysis revealed that across sessions, animals made more receptacle entries during the first 5 s of CS+ than in the CS− presentations (**Figure 2B**, main effect of stimulus and stimulus × session interaction, both *p* < 0.0001). In addition, the latency to first receptacle entry following CS+ onset decreased over subsequent conditioning sessions, reaching a plateau at 3-4 s (**Figure 2C**, main effect of session, *p* < 0.0001). Overall, the results indicate that both the control animals and the NR1^DATCreERT2^ mice learned the predictive value of the reward-paired stimulus and that they anticipated reward delivery during the CS+ presentation.

**Figure 1.**
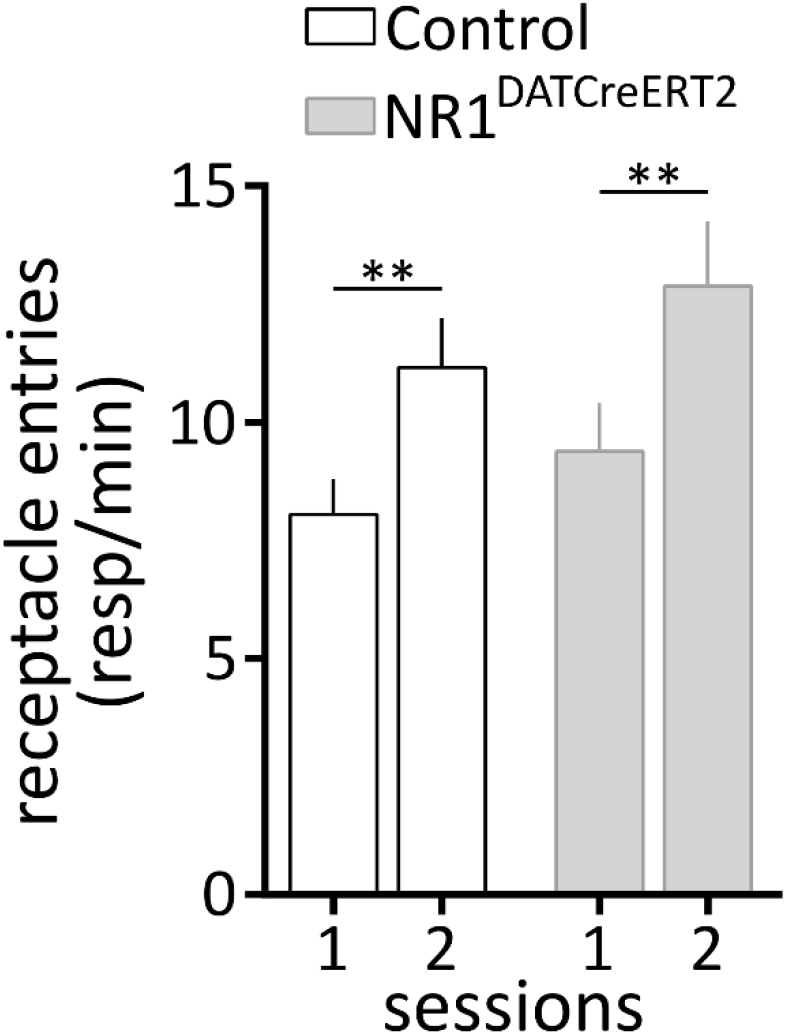
Adaptation to the water reward. The figure shows the average rate of water receptacle entries made by the control and the NR1^DATCreERT2^ mice during two consecutive days of adaptation to unconditioned reward delivery (n = 12 per group). Data are expressed as the mean ± SEM. ** *p* < 0.01, Bonferroni corrected t-test.

**Table 1.** Statistical table.

| Figure | Measure | Analysis | Effect | Statistical value | *p* value |
| --- | --- | --- | --- | --- | --- |
| 1 | Response rate | 2 way RM* ANOVA | session | $F_{1, 22} = 25.64$ | <0.0001 |
| | | | genotype | $F_{1, 22} = 1.37$ | 0.2539 |
| | | | session × genotype | $F_{1, 22} = 0.09$ | 0.7682 |
| 2A | Probability | mixed model ANOVA | stimulus | $F_{1, 22} = 128.33$ | <0.0001 |
| | | | session | $F_{10, 220} = 3.07$ | 0.0011 |
| | | | genotype | $F_{1, 22} = 1.41$ | 0.248 |
| | | | stimulus × session | $F_{10, 220} = 4.84$ | <0.0001 |
| | | | stimulus × genotype | $F_{1, 22} = 0.13$ | 0.725 |
| | | | session × genotype | $F_{10, 220} = 1.55$ | 0.1238 |
| | | | stimulus × session × genotype | $F_{10, 220} = 0.61$ | 0.803 |
| 2B | Receptacle entries | mixed model ANOVA | stimulus | $F_{1, 22} = 45.07$ | <0.0001 |
| | | | session | $F_{10, 220} = 0.52$ | 0.8773 |
| | | | genotype | $F_{1, 22} = 4.6$ | 0.0433 |
| | | | stimulus × session | $F_{10, 220} = 4.38$ | <0.0001 |
| | | | stimulus × genotype | $F_{1, 22} = 1.44$ | 0.244 |
| | | | session × genotype | $F_{10, 220} = 3.75$ | 0.0001 |
| | | | stimulus × session × genotype | $F_{10, 220} = 1.96$ | 0.0392 |
| 2C | Latency | 2 way RM ANOVA | session | $F_{10, 220} = 12.21$ | <0.0001 |
| | | | genotype | $F_{1, 22} = 0.96$ | 0.3369 |
| | | | session × genotype | $F_{10, 220} = 0.77$ | 0.6613 |
| 3 | Nose-poke responses | 2 way ANOVA | stimulus | $F_{1, 42} = 7.11$ | 0.0109 |
| | | | genotype | $F_{1, 42} = 7.34$ | 0.0097 |
| | | | stimulus × genotype | $F_{1, 42} = 0.55$ | 0.4645 |

\*RM – repeated measures

Although the mutation did not disrupt the ability to predict a reward, a reduced number of receptacle entries during CS+ trials in subsequent sessions was observed in the NR1^DATCreERT2^ mice (**Figure 2B**, session × genotype interaction, *p* < 0.001, and stimulus × session × genotype interaction, *p* < 0.05). This might indicate that over the course of conditioning, the CS+ acquired the incentive motivational properties that energized reward-seeking behavior in the control animals but not in the NR1^DATCreERT2^ mice. To further test whether the attribution of incentive value to the CS+ was impaired by the mutation, we performed a test for conditioned reinforcement. In this test, the water receptacle was replaced with two nose-poke ports. Making a nose-poke response into one of the ports resulted in CS+ presentation, while a nose-poke response made into the other port led to CS− presentation. Both groups preferentially made CS+ paired nose-poke responses, compared to CS− paired responses (**Figure 3**, main effect of stimulus, *p* = 0.01). However, the NR1^DATCreERT2^ mice made significantly fewer CS+ paired responses than the control animals (**Figure 3**, main effect of genotype, *p* = 0.01), indicating that for the mutant mice, the CS+ had a significantly weaker incentive value. This result confirms that the attribution of incentive salience to the reward-paired CS was impaired in the NR1^DATCreERT2^ mice.

**Figure 2.**
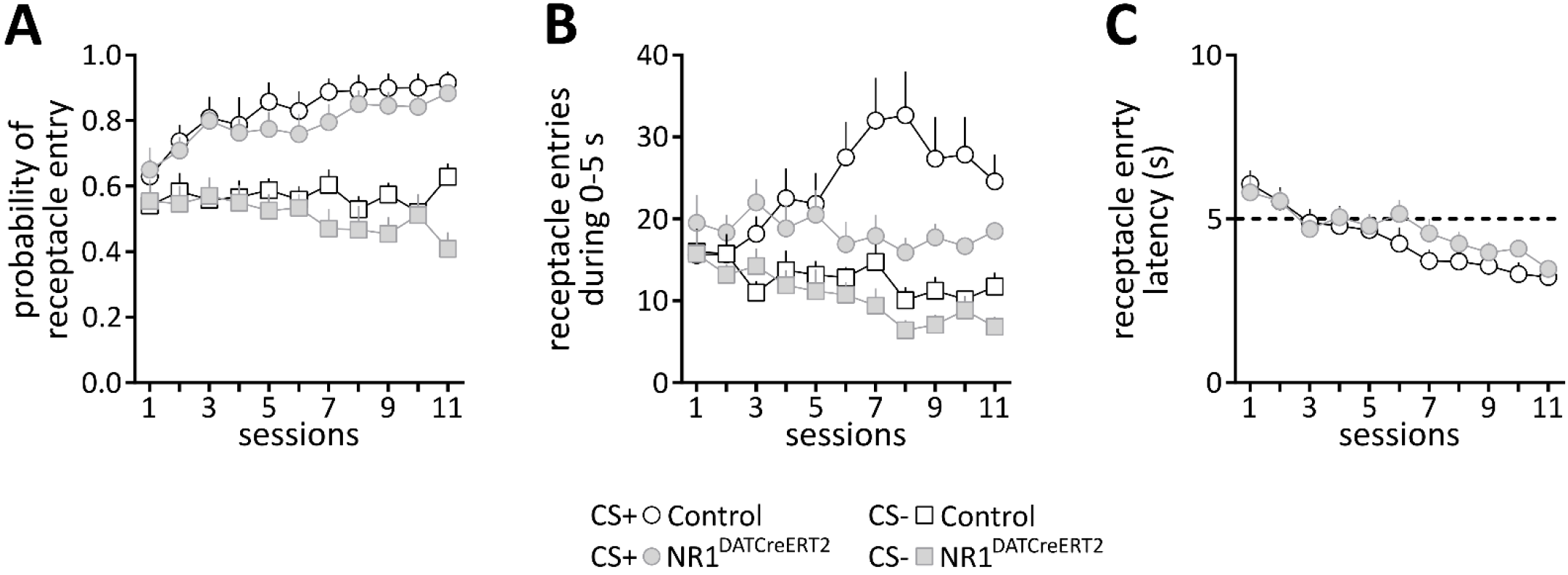
Stimulus-reward learning. (A) shows the probability of making an approach towards the water receptacle in CS+ and CS− trials. (B) shows the average number of receptacle entries made during the first 5 s from CS onset (i.e., before reward delivery in case of CS+ trials). (C) shows the time that elapsed from CS+ onset until the first visit to the water receptacle. Both groups n = 12. Data are expressed as the mean ± SEM.

**Figure 3.**
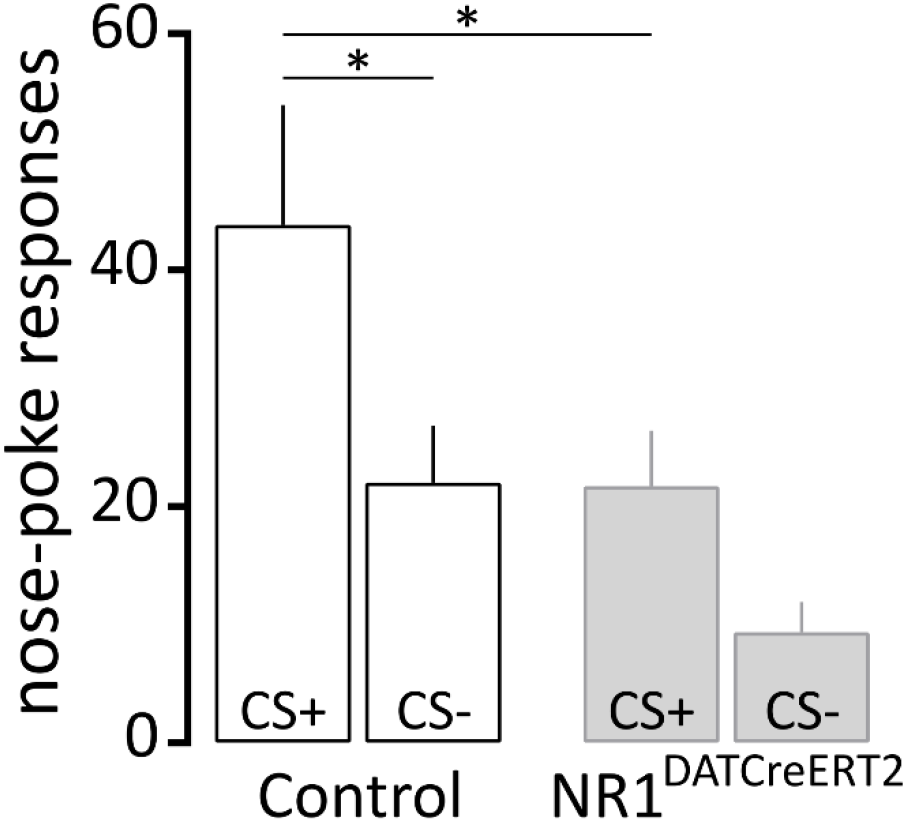
Conditioned reinforcement test. The figure shows the average number of nose-poke responses made by the control animals (n = 12) and the NR1^DATCreERT2^ mice (n = 11) in ports paired with CS+ or CS− presentations. One mutant animal was excluded from the analysis because no responses were made during the test and it died on the same day. Data are expressed as the mean ± SEM. * *p* < 0.05, Bonferroni corrected t-test.

Taken together, the results show that NMDA receptor-dependent signaling in DA neurons was not necessary for learning the predictive value of conditioned stimuli but was critical for the acquisition of incentive motivational properties by the reward-paired CS. The observation that reward prediction learning is preserved despite the absence of NMDA receptors in DA neurons is consistent with previous findings showing that mice with constitutive mutations also developed a conditioned response to a reward-paired cue (albeit in a slightly different paradigm) **[16, 17]**. Considering the proposed role of phasic DA signaling in reward prediction error coding, the behavior observed in the NR1^DATCreERT2^ mice is unexpected to an extent. However, the behavior is consistent with studies showing that DA mediates motivation but not reward learning **[20, 21]**. This indicates that our results are more consistent with the notion that DA signaling acts selectively in a form of stimulus-reward learning in which incentive salience is assigned to reward cues **[4]**. In this form of learning, the incentive properties acquired by a reward-paired stimulus can enable the stimulus to attract attention, energize reward-seeking or directly reinforce behavior. In agreement with this, we observed that the CS+ was able to energize reward-seeking behavior and to act as a conditioned reinforcer in control mice but not in mice with inducible ablation of NMDA receptors in DA neurons. This observation is in line with pharmacological studies showing that application of NMDA receptor antagonists into the VTA attenuated the expression of a conditioned approach to the reward receptacle during CS+ presentation and attenuated conditioned reinforcement **[10, 22]**. Nevertheless, our results differ from previous studies utilizing mouse models with constitutive NMDA receptor ablation. In particular, James and colleagues **[17]** tested the effects of the loss of NMDA receptors in DA neurons on incentive learning by focusing on sign-tracking behavior, a conditioned response directed towards reward-predicting stimulus. They found no effects of the mutation on sign-tracking, leading to the overall conclusion that NMDA receptor-dependent signaling in DA neurons is not involved in incentive learning. However, the authors noted that only a small fraction of mice showed sign-tracking behavior, suggesting that the analysis of sign-tracking may be less well suited for assessing incentive learning in mice.

Finally, it is worth noting that aberrant incentive learning has been long considered to play a role in the development of compulsive behaviors and addiction. Drug cues attributed with incentive salience can acquire the ability to control motivated behavior, triggering reinstatement of drug-seeking in animal models or relapse in humans. In line with this, NMDA receptor-dependent signaling in DA neurons was shown to be important for controlling relapse-like behavior. It was observed that the NR1^DATCreERT2^ mice were able to acquire a cocaine-induced conditioned place preference (CPP) but showed no reinstatement of CPP after a period of extinction **[15]**. This is consistent with our present findings, as the NR1^DATCreERT2^ mice were able to associate the reward-paired context (CS+) with the reinforcing effects of cocaine (US), but they failed to show reinstatement, likely because the CS+ was not attributed with incentive salience and after extinction had neither predictive nor motivational value. Moreover, it was reported that following a stable period of cocaine self-administration, the NR1^DATCreERT2^ mice showed reduced cue-induced reinstatement of cocaine-seeking behavior after forced withdrawal **[23]**. This suggested a reduced incentive motivation to engage in drug-seeking behavior in response to drug-paired cues. Overall, this means that NMDA receptors in DA neurons can regulate the attribution of incentive salience to stimuli associated with both natural rewards and drugs of abuse.

In conclusion, we showed that NMDA receptors play a crucial role in the attribution of incentive motivational value to reward paired cues. The results indicate that reward prediction learning does occur in the absence of key receptors implicated in the plasticity and activity of DA neurons, but the ability of reward-paired cues to invigorate and reinforce behavior is lost.

## Authors’ contributions

PEC designed and performed research. PEC and JRP analyzed the data and wrote the paper.

